# RhoGAP RGA-8 supports morphogenesis in *C. elegans* by polarizing epithelia through CDC-42

**DOI:** 10.1101/2019.12.15.877332

**Authors:** Hamidah Raduwan, Shashikala Sasidharan, Luigy Cordova Burgos, Andre G. Wallace, Martha C. Soto

## Abstract

CDC-42 regulation of non-muscle myosin/NMY-2 is required for polarity maintenance in the one-cell embryo of *C. elegans*. CDC-42 and NMY-2 regulate polarity throughout embryogenesis, but their contribution to later events of morphogenesis are less understood. We have shown that epidermal enclosure requires the GTPase CED-10/Rac1 and WAVE/Scar complex, its effector, to promote protrusions that drive enclosure through the branch actin regulator Arp2/3. Our analysis here of RGA-8, a homolog of SH3BP1/Rich1/ARHGAP17/Nadrin, with BAR and RhoGAP motifs, suggests it regulates CDC-42, so that NMY-2 promotes two events of epidermal morphogenesis: ventral enclosure and elongation. Genetic and molecular data suggest RGA-8 regulates CDC-42, and the CDC-42 effectors WSP-1 and MRCK-1, in parallel to F-BAR proteins TOCA-1 and TOCA-2. The RGA-8-CDC-42-WSP-1 pathway enriches myosin in migrating epidermal cells during ventral enclosure. We propose TOCA proteins and RGA-8 use BAR domains to localize and regenerate CDC-42 activity, thus regulating F-actin levels, through the branched actin regulator WSP-1, and myosin polarity through the myosin kinase MRCK-1. Regulated CDC-42 thus polarizes epithelia, to control cell migrations and cell shape changes of embryonic morphogenesis.

**Summary:** RGA-8, a protein with membrane binding and actin regulatory motifs, promotes embryonic morphogenesis by localizing active CDC-42 in developing epithelia, thus controlling actin and actin motors during cell movements.

## Introduction

Organ and tissue formation are highly regulated processes during embryonic development. During embryogenesis, epithelial cells must develop and maintain apicobasal polarity and healthy cell-cell junctions as they move past or over other tissues in the process of morphogenesis. Defects in this process can lead to birth defects, or premature death. Epidermal morphogenesis in the nematode *Caenorhabditis elegans* is an ideal model to study tissue morphogenesis ([1]Sulston et al, 1983; [2]Priess and Hirsh, 1986; [3]Chisholm and Hardin 2005).

Epidermal morphogenesis in *C. elegans* can be divided into several stages. Epidermal cells are born at the posterior and dorsal surface of the embryo and become arranged in three types of adjacent cells with distinct behaviors – two rows of dorsal cells, and on each side of the embryo, two rows of lateral seam cells, and two rows of ventral cells, (Fig. 1A). Epidermal morphogenesis begins when two outer rows, right and left ventral epidermal cells, migrate ventrally, pulling along other epidermal rows to enclose the embryo in a process known as ventral enclosure. This tissue migration is led by the two anterior-most ventral cells on each side, the leading cells. The leading cells reach the ventral midline first, while the more posterior ventral cells, the pocket cells, undergo a purse-like constriction, to enclose the embryo on the ventral side. Simultaneously, the two dorsal rows undergo dorsal intercalation to generate a single row of cells, in a process analogous to vertebrate convergent extension. Epidermal elongation, the process that squeezes the embryo into a worm shape, begins when the two rows of lateral seam cells increase their length along the anterior to posterior axis, and decrease along the dorsal/ventral axis. The dynamics of actin and actomyosin contractility that regulate epidermal morphogenesis are regulated by the Rho GTPases.

**Fig. 1.**
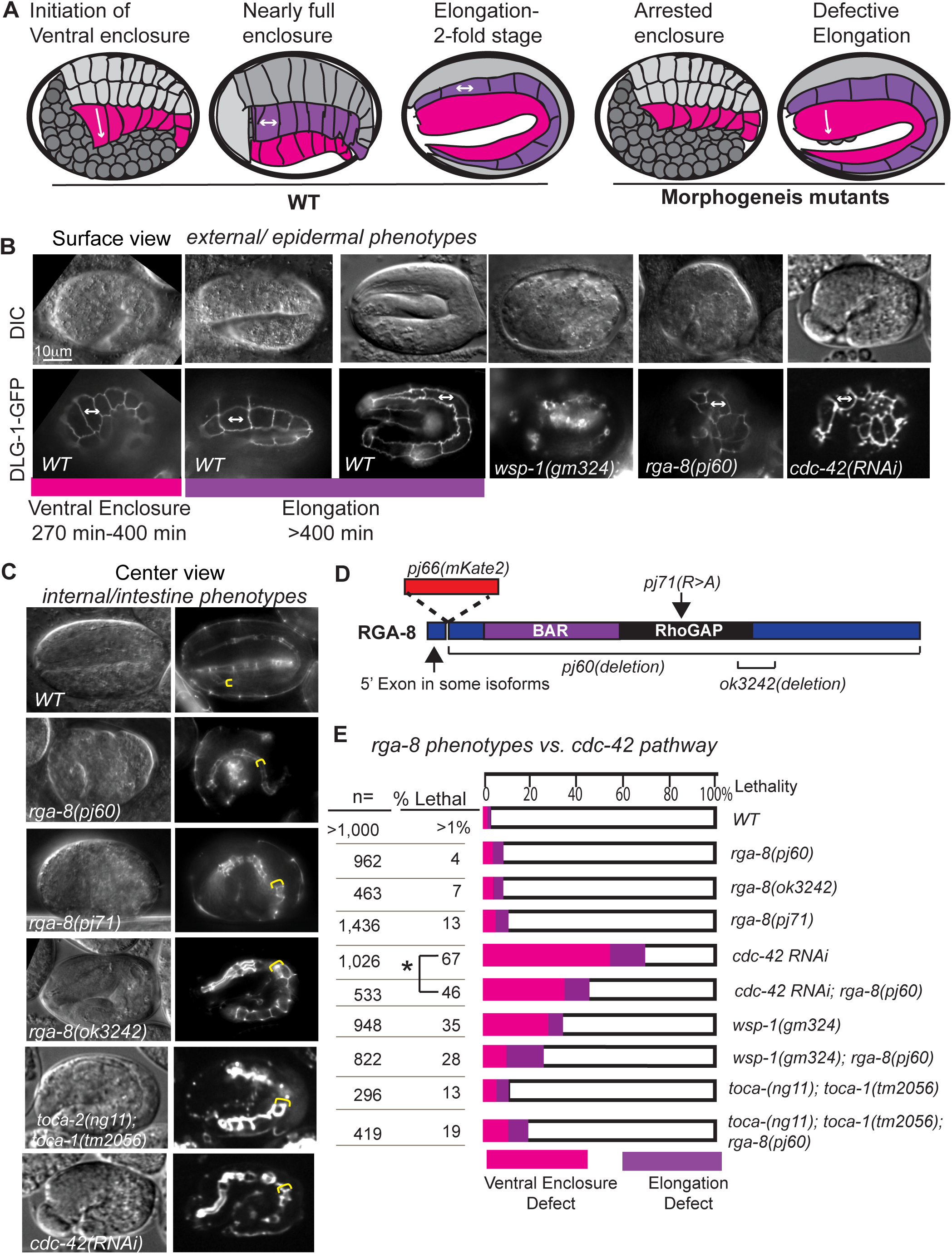
RhoGAP RGA-8 and CDC-42 pathway genes regulate two stages of morphogenesis. (A) Epidermal morphogenesis in *C. elegans* requires regulated movements of the cells of the ventral (magenta) and lateral (purple) rows. Ventral enclosure requires ventral-ward migrations (magenta) while during elongation shape changes in the lateral seam cells (purple) increase anterior-posterior length. Morphogenesis mutants arrest at either or both stages. (B) Wild type embryos shown beginning at 270 minutes after first cleavage, by differential interference contrast (DIC, top), and *dlg-1::gfp* (bottom), with a focus on the seam cells, illustrate cell shape changes of elongation. Mutant embryos are shown at the same time and focal plane. (C) Wild type and mutant embryos at 390 minutes after first cleavage, with a focus on the internal epithelia, illustrate elongation failures and widening of the apical intestine in the mutants. Yellow bracket: anterior end of intestine. (D) RGA-8 molecular model illustrates conserved BAR and RhoGAP domains. Genetic mutations from the *C. elegans* Gene Knockout Consortium (*ok3242* deletion) or from our CRISPR studies are indicated: *pj60* deletion, *pj71* GAP mutant and pj66 endogenous CRISPR N-terminal GFP tag, OX681 *rga-8(pj66)[mKate2::rga-8*] (see Fig. 2,3,4). (E) Comparison of the effects of *rga-8* mutations with mutations in *cdc-42* pathway genes. The numbers are taken from Table 1 and 2. In this Figure and others embryos are shown anterior to the left, and dorsal up, unless otherwise stated. Scale bars in all figures are 10um long. Embryonic times represent minutes after first cleavage at 23 C. Embryos were cultured and imaged at 23 C unless otherwise stated. Statistical significance in D was determined by Welch**’**s, or unequal variance, t-test. *P=<0.05.

In *C. elegans*, the three main GTPases, Rac1/CED-10, CDC-42 and RhoA/RHO-1, have been shown to be involved in some aspects of epidermal morphogenesis. Ventral enclosure is regulated by the GTPase Rac1/CED-10, which activates WAVE/Scar, a nucleation promoting factor (NPF) that activates branched actin formation through Arp2/3 [4,5,6]. Rac1/CED-10 and WAVE/Scar also regulate dorsal intercalation ([7]Soto et al., 2002; [8]Patel et al., 2008; [9]Walck-Shannon et al., 2015), underscoring the important role played by CED-10/Rac1 in regulating the overall process of epidermal morphogenesis. In parallel to the CED-10/Rac1/WAVE pathway, morphogenesis is supported by the CDC-42 GTPase and its Nucleation Promoting Factor, WASP ([10]Sawa et al., 2003; [11]Withee et al., 2004; [12]Walck-Shannon et al., 2016).

**Table 1:**
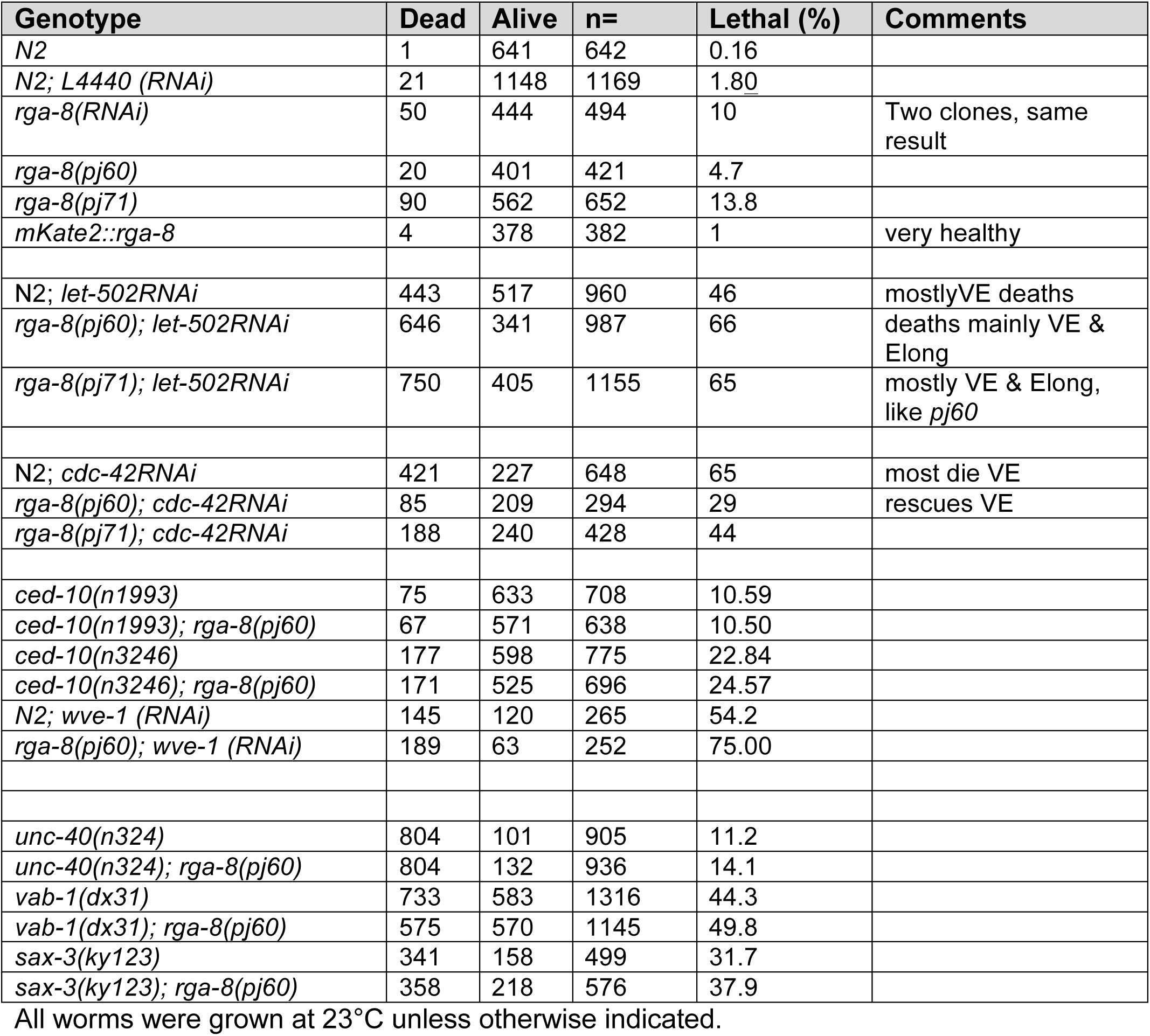
Embryonic lethality of *rga-8* alleles, vs. regulators of embryonic F-actin.

**Table 2:** *rga-8* and *cdc-42* pathway phenotypes sorted by time of arrest.

| **Lethal Phenotypes\* (percent).....** |  |  |  |  |  |
| --- | --- | --- | --- | --- | --- |
| **Genotype** | **Early arrest** | **Ventral Enclose** | **Elongation** | **Total Lethal** | **n=** |
| *rga-8(pj60)* | 0.5% | 1.6% | 1.3% | 3.4% | 561 |
| *rga-8(pj71)* | 0.5 | 7.9 | 4.6 | 13 | 784 |
| *wsp-1(gm324)* | 2.9 | 28 | 4 | 35 | 948 |
| *toca-2; toca-1* | 0.3 | 4.4 | 7.8 | 12.5 | 888 |
| *rga-8(pj60);wsp-1(gm324)* | 0.4 | 9.7 | 11 | 21.3 | 822 |
| *rga-8(pj60); toca-2* | 0.5 | 12.4 | 5.7 | 18.6 | 419 |
| *dlg-1::gfp* | 0 | 0.4 | 0.4 | 0.9 | 1164 |
| *dlg-1::gfp; rga-8(pj71)* | 0.15 | 6.3 | 4.1 | 10.6 | 678 |
| *dlg-1::gfp; rga-8(ok3246)* | 0.5 | 2.8 | 5.2 | 8.4 | 758 |
| *dlg-1::gfp; wsp-1(gm324)* | 1.9 | 20.2 | 7.2 | 29 | 1135 |
| *dlg-1::gfp; toca-2; toca-1* | 0.9 | 18 | 17 | 36 | 438 |
| *dlg-1::gfp; toca-2; toca-1; rga-8(pj60)* | 1.6 | 10.6 | 6.5 | 19 | 904 |
| *rga-8-RNAi* | 0.3 | 1.7 | 4.4 | 6.4 | 939 |
| *wsp-1(gm324); rga-8-RNAi* | 2.2 | 22.3 | 6.3 | 30.7 | 557 |
| *toca-2; toca-1; rga-8-RNAi* | 1.0 | 19.2 | 8.5 | 29 | 273 |
| *mrck-1 RNAi* | 0 | 7.6 | 12.3 | 20 | 236 |
| *pj60; mrck-1 RNAi* | 0.3 | 10.8 | 15.7 | 27 | 655 |
| *cdc-42 RNAi Day 2* | 2.6 | 61 | 10 | 71 | 378 |
| *cdc-42 RNAi; rga-8(pj60)* | 0 | 49 | 19 | 68 | 239 |
| *cdc-42 RNAi; rga-8(pj71)* | 0 | 37 | 24 | 61 | 268 |
| *cdc-42 RNAi; toca-2; toca-1* | 2 | 53 | 24 | 78 | 152 |

\*Explanation of the phenotypes:
*Early arrest*: includes embryos with cytokinesis defects and other early arrest so that tissue differentiation is defective. Lethal arrest occurs before epidermal morphogenesis begins.
*Ventral enclosure*: describes embryos with fully differentiated tissues that fail epidermal enclosure so that internal organs end up on the outside (AKA Gex, gut on exterior).
*Elongation*: describes embryos that enclose but then fail to lengthen along the anterior/posterior axis. Some of these burst after initially enclosing.

Epidermal elongation depends on regulated actomyosin driven by the GTPase RHO-1/RhoA (Reviewed in [13]Vuong-Brender et al., 2016). Actomyosin is negatively regulated by MEL-11/MYPT, which is expressed in the dorsal and ventral epidermis ([13]Wissman et al., 1999). In the absence of MEL-11 embryos burst due to increased tension on adherens junctions. Actomyosin contractility in the seam cells is promoted by Rho Kinase LET-502/ROCK, which is expressed in the epidermal seam cells ([13]Wissman et al., 1999). Removing LET-502/ROCK caused defects in elongation of the embryo. Activation of myosin II is achieved mainly through the LET-502/Rho kinase, but two additional kinases can contribute to maintain myosin II activity. P21-activated kinase PAK-1 and CDC-42-activated kinase MRCK-1 act in parallel to LET-502, since their loss enhances *let-502* mutant severity. MRCK-1 acts upstream of MEL-11 ([14]Gally et al., 2009) and is under the regulation of CDC-42 [15].

Studies of *cdc-42* function in *C. elegans* have mostly focused on its role during the one-cell stage, where it helps set up anterior/posterior polarity as a component of the anterior PAR complex (Reviewed in [16]Cowan and Hyman 2007). CDC-42 in the one cell embryo also regulates myosin levels and cortical enrichment ([17]Kumfer et al., 2010; [18]Motegi and Sugimoto, 2006; [19]Schonegg and Hyman, 2006; [20]Beatty et al., 2013). Studying the morphogenesis role requires getting past the early roles for CDC-42 in setting up polarity. A role for CDC-42 and WASP/WSP-1 during epidermal morphogenesis has been proposed ([20]Sawa et al, 2003; [11]Withee et al., 2004). Oulette and colleagues found that overexpression of CDC-42 caused altered protrusions of leading cells, and embryonic lethality, suggesting a balance of active and inactive CDC-42 is required for efficient ventral migration ([21]Ouellette et al., 2016). Transducer of cytokinesis-1, TOCA-1, contains a GTPase binding domain (GBD) that binds to CDC-42, an SH3 domain that binds to proline-rich region of WASP or other proteins, and an F-BAR domain that senses and binds to curved membranes ([22]Takano et al., 2008). *C. elegans* has two partially redundant homologs of TOCA1, named TOCA-1 and TOCA-2, which were both implicated in epidermal morphogenesis of *C. elegans* ([23]Giuliani et al., 2009). The Rho GAP RGA-7 was shown to genetically interact with TOCA-1/2 and CDC-42 during the formation of actin-rich protrusion in leading cells ([21]Ouellette et al., 2016). These results suggested that the conserved pathway of branched actin regulation by TOCA-1/2-CDC-42-WASP also regulates ventral enclosure during *C. elegans* morphogenesis. Zilberman and colleagues used ZIF1 degron technology to partially rescue *cdc-42* null mutants to survive past the one-cell defects to study epidermal morphogenesis, and uncovered ventral enclosure and elongation defects in *cdc-42* mutants ([24]Zilberman et al., 2017). CDC-42 also has a role during dorsal intercalation ([12]Walck-Shannon et al., 2016). These reports suggest an overall role for CDC-42 during epidermal morphogenesis.

The role actomyosin contractility plays during ventral migration is not clear. ANI-1, a multidomain protein that organizes actomyosin contractility, is required to ensure proper alignment of contralateral leading-edge epidermal cells meeting at the ventral midline. However, expression of ANI-1 was not detected in the epidermal cells, leading to the suggestion that ANI-1’s action may be due to the interaction between the underlying neuroblast (neuron precursors) with the migrating epidermal cells. Interestingly, loss of *rho-1, let-502* or *mel-11* led to ventral enclosure defects, but it was not clear if this was due to roles in the neuroblasts or in the epidermis ([25]Fotopoulos et al., 2013). Studies of HUM-7, a GAP that regulates epidermal morphogenesis through RHO-1/RhoA showed it affected NMY-2::GFP levels specifically in the migrating epidermal cells ([26]Wallace et al., 2018). Altogether, the evidence suggests RHO-1/RhoA and myosin also contribute to ventral enclosure.

Here we investigate a proposed novel regulator of CDC-42 during ventral enclosure in *C. elegans.* We identify the RhoGAP and BAR domain protein, RGA-8, as a regulator of CDC-42, based on genetic interactions with *cdc-42, wsp-1*, and with the *toca-2(ng11);toca-1(tm2056)* double mutant. A CRISPR generated *rga-8* null allele, *pj60*, is tested for effects on a CDC-42 biosensor. Using RGA-8 endogenously-tagged via CRISPR, we examine RGA-8 distribution in epidermal cells and in polarized epithelia of the pharynx and intestine. The *rga-8(pj60)* null mutant and WASP-1 mutant *wsp-1(gm324)* are examined for effects on non-muscle myosin II (NMY-2::GFP) at the leading edge of migrating epidermal pocket cells during ventral enclosure. We propose a model where RGA-8 and TOCA-1/TOCA-2, through CDC-42, WSP-1 and myosin kinase MRCK-1, enrich and maintain polarized NMY-2::GFP accumulation at the epidermal pocket cells to support ventral enclosure.

## Results

### RGA-8 is a GAP for CDC-42 that regulates *C. elegans* embryonic morphogenesis

To characterize the regulators of CDC-42 during epidermal morphogenesis, we screened the *C. elegans* family of GTPase activating proteins (GAPs), which inactivate and turn over the GTPases ([27]Neukomm et al., 2011) and compared them to the epidermal morphogenesis phenotype for CDC-42 pathway genes using *dlg-1::gfp* transgenic strain (FT250; [28]Totong et al., 2007). First we knocked down *cdc-42* via RNAi by feeding *cdc-42* double stranded RNAi to L1 worms and waiting 2 days. Under these conditions the RNAi knockdown resulted in approximately 70% embryonic lethality in their progeny, yielding the highest embryonic death during ventral enclosure stage at 61%, followed by elongation stage at 10%, and early differentiation at only 2.6% (Fig. 1 A-D, Table 2). We hypothesized that removing GAPs from *cdc-42* RNAi, which creates a hypomorphic *cdc-42* background, might rescue embryonic lethality. We focused on one candidate CDC-42 GAP, named Rho-GTPase activating protein-8 (RGA-8). We generated an *rga-8* deletion allele, *pj60*, using CRISPR (Fig. 1D) and found it rescued *cdc-42 RNAi* embryonic lethality from 67% to 46% (Fig. 1E). *rga-8* was thus a candidate GAP for CDC-42.

To investigate which step of embryogenesis is regulated by RGA-8 and the CDC-42 pathway, we crossed mutants or fed RNAi food to *dlg-1::gfp* which is expressed at *C. elegans* apical junctions, and thus allowed us to monitor cell migration and cell shape changes. *dlg-1::gfp; rga-8(pj60)*, resulted in 4% embryonic lethality, with approximately half the embryos arresting during ventral enclosure, and the other half during elongation (Tables 1,2). Another *rga-8* allele, *ok3242*, a 611 bp deletion that causes a frame shift and premature stop codon, truncating RGA-8 part way through the GAP domain (Fig. 1D), showed higher embryonic lethality, at 7% (8.4% in *dlg-1::gfp* background) (Fig. 1C,E). We inactivated the GAP domain of RGA-8 by creating a CRISPR allele, *pj71*, which switches the catalytic Arginine to an Alanine, and measured 13% embryonic lethality, with arrests during ventral enclosure and elongation (Fig. 1C,D,E). Thus, *rga-8* has a role in two steps of epidermal morphogenesis, and its RhoGAP domain appears to be involved in this process.

We next investigated if RGA-8 interacted with known components of CDC-42 pathway: TOCA-1, TOCA-2 and WASP/WSP-1. WASP/WSP-1 is a branched actin nucleation promoting factor that is autoinhibited until it binds TOCA and CDC-42. Activated WASP-1 promotes branched actin formation through the Arp-2/3 complex (Reviewed in [29]Kurisu & Takenawa 2009; [30]Takenawa & Suetsugu 2007). TOCAs are F-BAR domain proteins that recruit regulators of branched actin to subcellular domains. Deletion mutations *wsp-1(gm324)* and *toca-2(ng11);toca-1(tm2056)* led to embryonic lethality during ventral enclosure and elongation at 35% and 13% respectively, as previously reported (Table 2, Fig. 1, [11]Withee et al., 2004; [23]Giuliani et al., 2009). When the *rga-8(pj60)* deletion was crossed to the *toca-2(ng11);toca-1(tm2056)* double mutant, it enhanced embryonic lethality from 13% to 19% (Table 2). Genetic doubles with *wsp-1(gm324)*, a known effector of *cdc-42* that regulates ventral enclosure [5], resulted in embryonic lethality more similar to loss of *wsp-1*, at approximately 28%, suggesting that WSP-1 may be epistatic to RGA-8. Collectively, our results suggested that RGA-8 works with CDC-42 to regulate embryogenesis, possibly parallel to TOCA-1/TOCA-2, and upstream of WSP-1.

### RGA-8 acts in parallel to RHO-1 and CED-10/Rac1 during *C. elegans* morphogenesis

The homologs of RGA-8, SH3BP1, RICH-1 and NADRIN, are proposed GAPs for the GTPase Rac1. If RGA-8 was a GAP for the GTPase Rac1/CED-10, we expected it to rescue the hypomorphic alleles of *ced-10*. Genetic doubles with two hypomorphic alleles, *ced-10(n1993)* and *ced-10(n3246)*, resulted in lethality that reflected the *ced-10* alleles, from 10.6% to 10.5%, and 22.8% to 24.6% (Table 1). Combining the *pj60* mutant with partial depletion of the Rac-1-dependent WAVE/WVE-1 protein, via *wve-1 RNAi*, ([8]Patel et al., 2008), led to an enhancement of embryonic lethality, from 54.7% to 75% (Table 1). This suggested that RGA-8 is not a Rac1 GAP and instead functions in parallel with the Rac1-WAVE/SCAR branched actin pathway to mediate embryogenesis of *C. elegans.*

We tested RGA-8 genetic interactions with the RHO-1 pathway. Embryonic lethality due to RNAi of the Rho-1 kinase *let-502* was increased by the putative null allele *rga-8(pj60)* from 46% to 66%, and also increased by the *pj71* GAP mutant. Embryonic lethality due to reduced myosin phosphatase, *mel-11*, via *RNAi*, was increased in *rga-8(pj60)* or *rga-8(pj71)* embryos (Table 1). These results suggest *rga-8* acts in parallel to the RHO-1 pathway. Collectively, we suggest RGA-8 is a GAP for CDC-42, since putative null deletion mutations and RNAi depletion show genetic interactions with components of CDC-42 pathway signaling, including suppression of *cdc-42* lethality.

### RGA-8 is expressed in most cells with preferred localization at apical pharyngeal and intestinal cells

To examine the expression pattern of RGA-8, we endogenously-tagged it at the N-terminus with mKate2, a red fluorophore, using CRISPR technology ([31]Dickinson et al, 2015) (Fig. 1D). Confocal spinning disk microscopy showed diffuse signal in most cells, with enrichment of *mKate2::rga-8* at apical regions of two epithelia, pharynx and intestine (Fig. 2A,B). Apical enrichment starts at approximately 240 min after first cleavage, at a time when apical junctions are being established, and is maintained until adulthood. Apical enrichment in epithelia is also seen in the transgenic strain, *gfp::cdc-42* (Fig. 2A,B; [17]Kumfer et al., 2010).

**Fig. 2.**
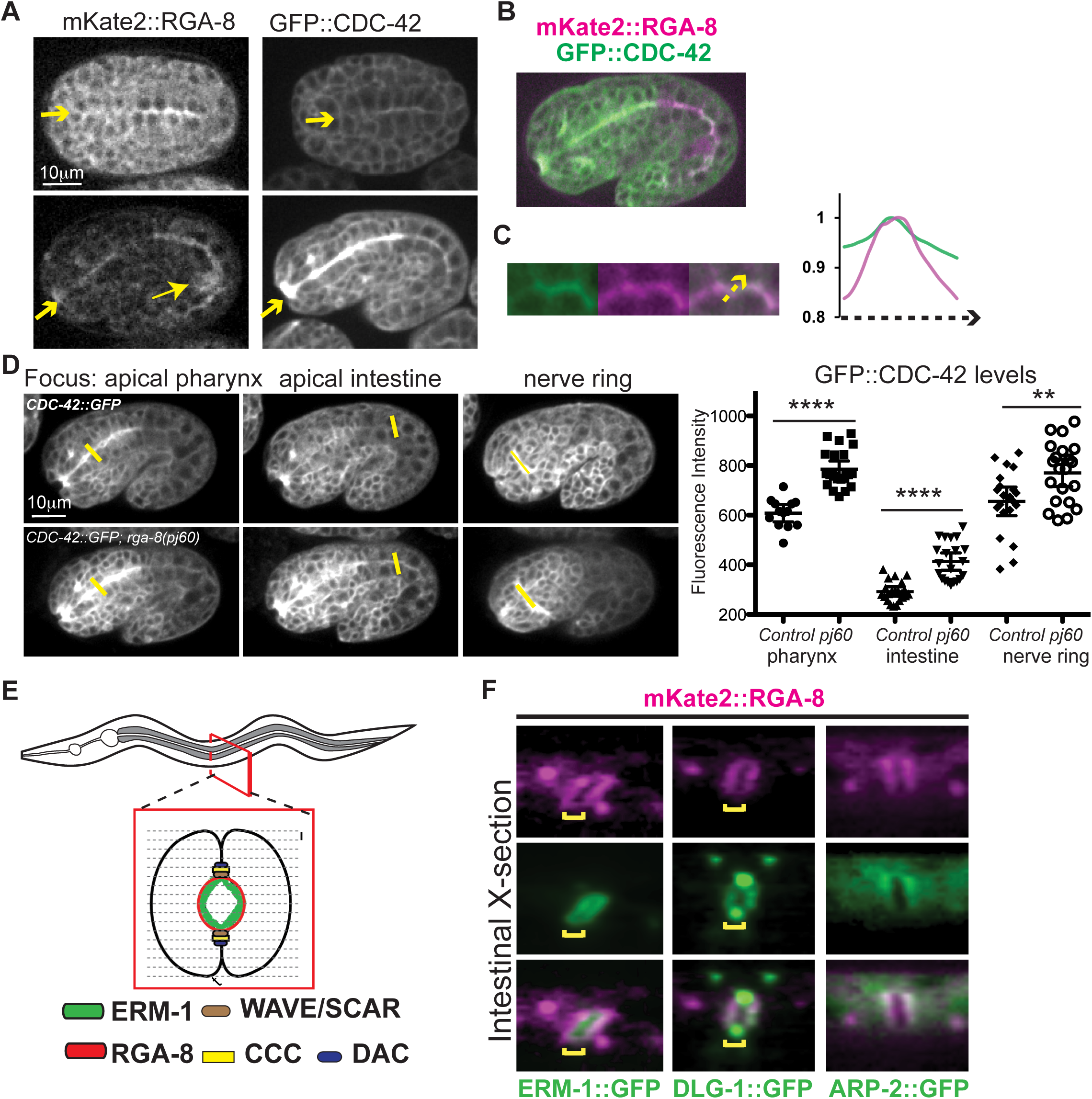
RGA-8 is enriched apically in pharyngeal and intestinal epithelia. (A-C) Embryos expressing *mKate2::rga-8*, or *gfp::cdc-42*, or both (color images). Embryos at 240 minutes, top row, (A), and 360 minutes, bottom, (A) and (B). Arrows at anterior point to apical pharynx. Arrow head, lower left, indicates bright signal in the two germline cells. (C), a crop from the intestine in B, includes a line scan to compare enrichment at the apical intestine. (D). Control *gfp::cdc-42* embryos were compared to *gfp::cdc-42; rga-8(pj60)* embryos for enrichment at apical pharynx, intestine and at the nerve ring. Different focal planes are shown. (E) Cartoon illustrating how images for (F) were generated. L1 larvae were imaged every 1μm across the entire worm. A cross-section reconstruction was generated by a line scan along all z-sections across the larval intestine. Cross sections of the intestine of L1 larvae compare mKate2::RGA-8 with ERM-1::GFP, DLG-1::GFP, and ARP-2::GFP. Statistical significance in E was determined by two tailed student’s t test with Welch’s correction. P Values **= <0.001, ****= < 0.0001.

#### RGA-8 regulates apical enrichment of GFP::CDC-42

If RGA-8 is a GAP that regulates CDC-42, it may alter the levels of CDC-42. Crossing the *rga-8(pj60)* deletion allele into a rescuing transgene, *gfp::cdc-42* (WS4700; [32]Neukomm et al., 2014), resulted in increased *gfp::cdc-42* levels at the apical intestine and apical pharynx (Fig. 2D). Other regions with high *gfp::cdc-42* expression, like the nerve ring, also showed higher expression in *rga-8(pj60).* This result suggested that RGA-8 regulates CDC-42.

#### Intestinal expression

We examined the subcellular localization of *mKate2::rga-8*, relative to other proteins, at the apical intestine in L1 larvae. Viewing the images as cross-section projections, showed *mKate2::rga-8* is enriched all around the lumen of the intestine (Fig. 2E,F). This is different than the localization of apical junction marker, *dlg-1::gfp*, that localizes at two puncta (Fig. 2F). We compared *mKate2::rga-8* to *erm-1::gfp* which is enriched at the most apical regions of intestine ([33]Gobel et al., 2004) and found that *mKate2::rga-8* localizes more basally than *erm-1::gfp*, suggesting that RGA-8 is not found in the microvilli Fig. 2F). In agreement with this, EM studies of *rga-8(pj60)* showed no dramatic changes at microvilli at the apical intestine in adults (data not shown). Comparing *mKate2::rga-8* to *arp-2::gfp* showed similar patterns all around the apical side of the intestinal lumen (Fig. 2F). We conclude that RGA-8 is enriched around the intestinal lumen, basal to the *erm-1::gfp* apical-most domain and partially overlaps with *arp-2::gfp* localization.

#### Epidermal expression

Since RGA-8 regulates ventral enclosure, we examined if it is expressed in the epidermis. Using an epidermal F-actin transgene, *plin-26::Lifeact::gfp* [34], we found that *mKate2::rga-8* is expressed diffusely in the epidermal cells, with some enrichment at cell membranes, similarly to regulators of branched actin, like ARP-2::GFP (Fig. 3A,B). *mKate2::rga-8* in epidermal cells localizes basally to two apical junction complexes as shown by comparison to the cadherin component HMP-1::GFP/alpha catenin, and the DLG-1/AJM-1 complex component, DLG-1::GFP (Fig. 3B). We conclude that RGA-8 is expressed in the epidermal cells, and it is enriched in regions that are also enriched with actin regulators.

**Fig. 3.**
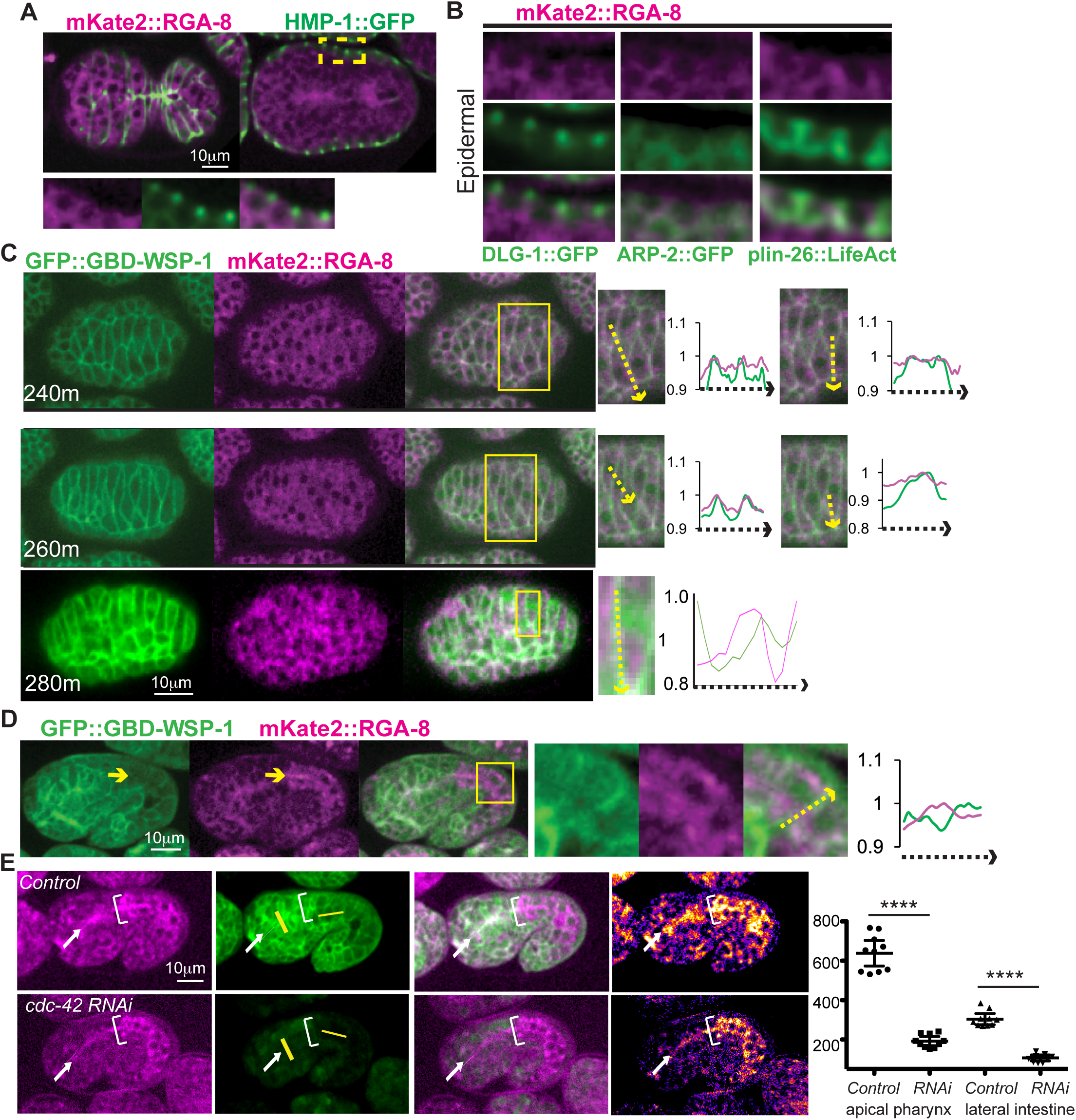
mKate2::RGA-8 localization relative to epidermally enriched proteins, including a biosensor for active CDC-42. (A) 300 minute embryo shown ventral up, surface focus (left) or internal focus (right) as epidermis meets on the ventral side, expressing *mKate2::rga-8* and *hmp-1::gfp* ([43]Marston et al., 2016) to illustrate epidermal apical epithelial junctions. Yellow square indicates region cropped to focus on epidermis. (B) Localization of *mKate2::rga-8* compared to *dlg-1::gfp, arp-2::gfp, Plin26::LifeAct::gfp* in crops made as in A. (C) Embryos shown dorsal up at approximately 240 and 260 minutes, during dorsal intercalation. *gbd-wsp-1::gfp* was shown to enrich in intercalating epidermis ([12]Walck-Shannon et al., 2016). *mKate2::rga-8* is shown relative to *gbd-wsp-1::gfp*. Line scans illustrate patterns of enrichment. (D) Embryos at 360 min, anterior left and dorsal up, focused on internal epithelia show localization of *mKate2::rga-8* relative to *gbd-wsp-1::gfp*. In crop of intestine a line scan (yellow arrow) compares apical enrichment. (E). *mKate2::rga-8*; *gbd-wsp-1::gfp* control embryos (top) or with *cdc-42* RNAi depletion (bottom). Right-most two panels show *mKate2::rga-8* in “Fire” mode, ImageJ, to better show differences in intensity. Graphs compare effects on *gbd-wsp-1::gfp* in apical pharynx or intestine. White arrow points to apical pharynx, white bracket shows anterior of the intestine, yellow lines show where measurements were made in apical pharynx and lateral intestine. Statistical significance in (E) was determined by Welch**’**s, or unequal variance, t-test. n= at least 10 embryos. P Values ****= < 0.0001.

### RGA-8 colocalizes with active CDC-42 in some tissues

#### Dorsal epidermis

CDC-42 was proposed to regulate dorsal intercalation, using a biosensor for active CDC-42 *gbd-wsp-1::gfp*, where the G protein-binding domain of WSP-1, (GBD)WSP-1, is fused to GFP ([17]Kumpfer, 2010, [12]Walck-Shannon, 2016). The same *gbd-wsp-1::gfp* domain, under the *cdc-42* promoter, shows specific localization to the cell-cell junctions of migrating epidermal cells ([24]Zilberman et al. 2017). We tested if RGA-8 colocalizes with active CDC-42 by crossing *mKate2::rga-8* into *Pcdc-42*::*gbd-wsp-1::gfp* (Fig. 3C). Line scans through the dorsal cells during intercalation showed *mKate2::rga-8* appears enriched at the cortex of intercalating cells, with transient medial enrichment, similar to *Pcdc-42*::*gbd-wsp-1::gfp* (Fig. 3C).

#### Loss of CDC-42 affects Pcdc-42::gbd-wsp-1::gfp but not mKate::rga-8 at apical intestine

If RGA-8 is a GAP for CDC-42, we predicted loss of CDC-42 would not affect it. While *mKate2::rga-8* and *gfp::cdc-42* co-localize at both the apical pharynx and intestinal (Fig. 2A) *mKate2::rga-8* and *Pcdc-42*::*gbd-wsp-1::gfp* only colocalize at the apical pharynx. In the intestine *Pcdc-42*::*gbd-wsp-1::gfp* is instead enriched basolaterally and seems depleted apically (Fig. 3D). Loss of *cdc-42* via RNAi resulted in strong depletion of *Pcdc-42*::*gbd-wsp-1::gfp*, as expected, including loss of the robust apical pharynx enrichment. In contrast, the localization of *mKate2::rga-8* was maintained, including robust apical enrichment in the pharynx (Fig. 3E). This result suggests that RGA-8 is not regulated by CDC-42.

### RGA-8 regulates apical/basal distribution of a CDC-42 biosensor

If RGA-8 is a CDC-42 GAP, it may be responsible for the apical/basal distribution of active CDC-42/GBD-WSP-1::GFP. In support of this, crossing the *rga-8* null allele *pj60* into *gbd-wsp-1::gfp* resulted in a significant change in apical enrichment of GBD-WSP-1::GFP in the pharynx (Fig. 4A). Control embryos showed higher apical to basal GBD-WSP-1::GFP, (1.45 fold higher) while *pj60* embryos showed reduced overall signal, and reduced apical to basal distribution (1.15 fold). In the intestine, where control embryos are instead basolaterally enriched, *pj60* embryos show decreased basolateral membrane signal. The lateral to basal ratio dropped from 1.1 to 0.7 (Fig. 4B). While overall cytoplasmic and membrane signals are decreased in *pj60* embryos, the changes in both tissues suggest an apical/basal shift of active CDC-42.

**Fig. 4.**
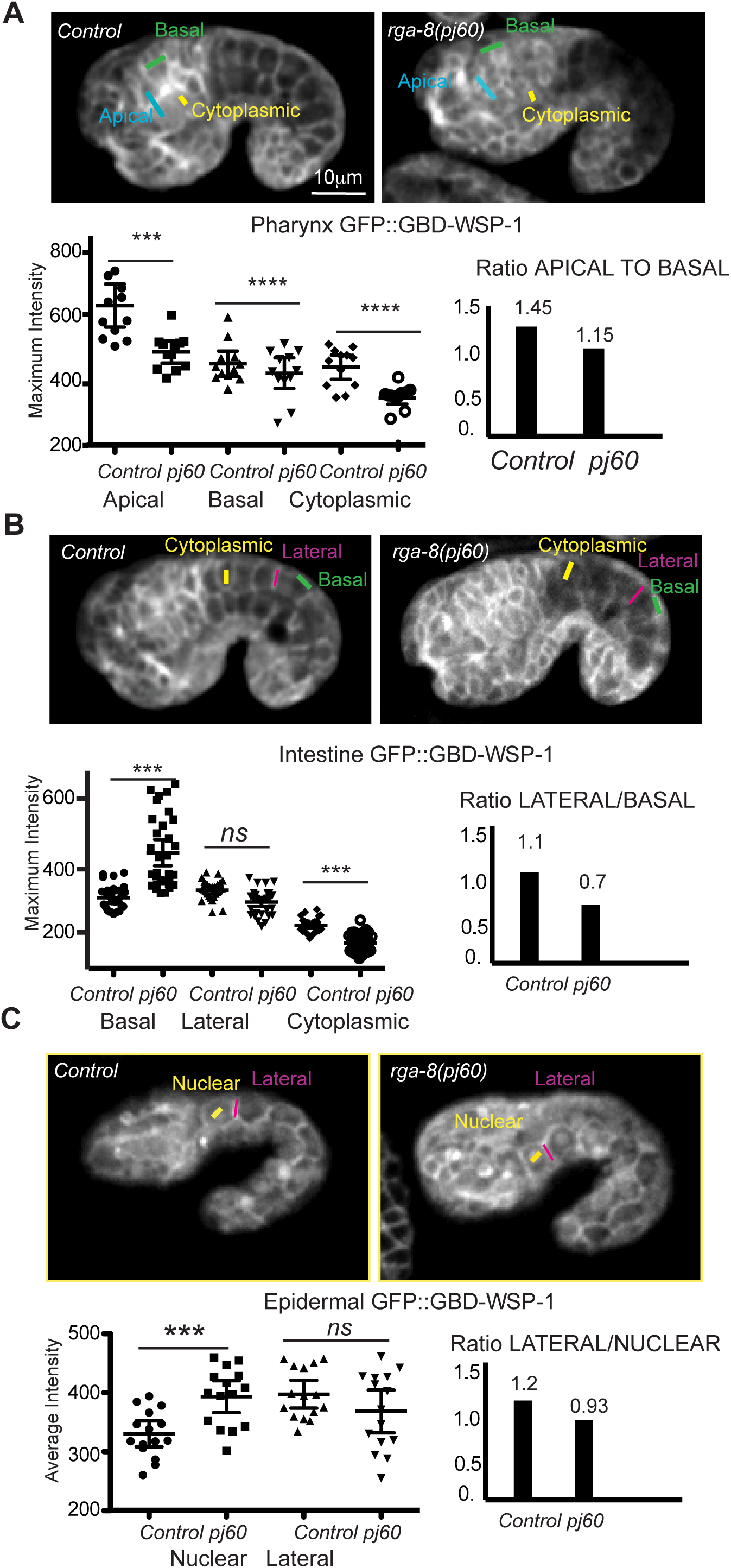
RGA-8 regulates apical/basal distribution of active CDC-42 in two epithelia. Distribution of *gbd-wsp-1::gfp* in control embryos and in *rga-8(pj60)* mutant embryos. (A) 360 minute embryos, shown with focus on the apical pharynx. (B). 360 minute embryos, shown with focus on the apical intestine. (C) 390 minute embryos, focus on the lateral seam cells. n= at least 7 embryos per type of measurements, and 4 cells were measured per embryo. Regions measured are indicated on each image. For A & B, measurements are maximum intensities. For C, measurements are averages. Statistical significance was determined by two-tailed student’s t test with Welch’s correction. P Values ***= <0.001, ****= < 0.0001.

#### RGA-8 localization relative to CDC-42 biosensor in epidermis

To test if RGA-8 regulates active CDC-42 in the epidermis, we first examined the *gbd-wsp-1::gfp* pattern at the ventral epidermis during enclosure. *gbd-wsp-1::gfp* is enriched at cell boundaries, but we did not detect obvious enrichment at the ventral midline. We examined the lateral seam cells that drive the first part of epidermal elongation. *Pcdc-42*::*gbd-wsp-1::gfp* is normally enriched at all boundaries of the seam cells and loss of *cdc-42* in this strain results in decreased membrane signal, and increased nuclear signal (Fig. 3C; [24]Zilberman et al., 2017). In *Pcdc-42*::*gbd-wsp-1::gfp*; *rga-8(pj60)* embryos there was a slight drop in the membrane signal at lateral regions of the seam cells and increased nuclear signal. While the lateral decrease was not significant, the nuclear increase was, and the ratio of lateral to nuclear signal decreased (Fig. 4C). Altogether, our results show RGA-8 promotes polarized distribution of a CDC-42 biosensor at membranes.

### Defects in CDC-42 pathway alter F-actin levels in migrating epidermis

The Rac/WAVE/Scar pathway is known to regulate F-actin levels, polarity and dynamics in migrating epidermal cells ([35]Bernadskaya et al., 2012), but the effects of the CDC-42 pathway on F-actin in migrating epidermis is less understood. We monitored levels of epidermal F-actin by crossing genetic mutants into *plin-26::LifeAct::GFP* or *plin-26::LifeAct::mCherry*, ([36]Havrylenko et al., 2015) depending on the chromosomes where the genes are found. We detected increased F-actin in *wsp-1(gm324)* and *rga-8(pj60)* mutants in leading-edge cells undergoing ventral enclosure, relative to controls. Similarly, depleting *cdc-42* via RNAi, and measuring F-actin only in embryos that do not arrest early, showed elevated epidermal F-actin levels (Fig. 5A, B). *toca-2; toca-1* double mutants did not significantly alter epidermal F-actin levels. Therefore, a proposed CDC-42 pathway that includes RGA-8 and WSP-1 appears to be required to maintain appropriate F-actin levels in the migrating epidermal cells.

**Fig. 5.**
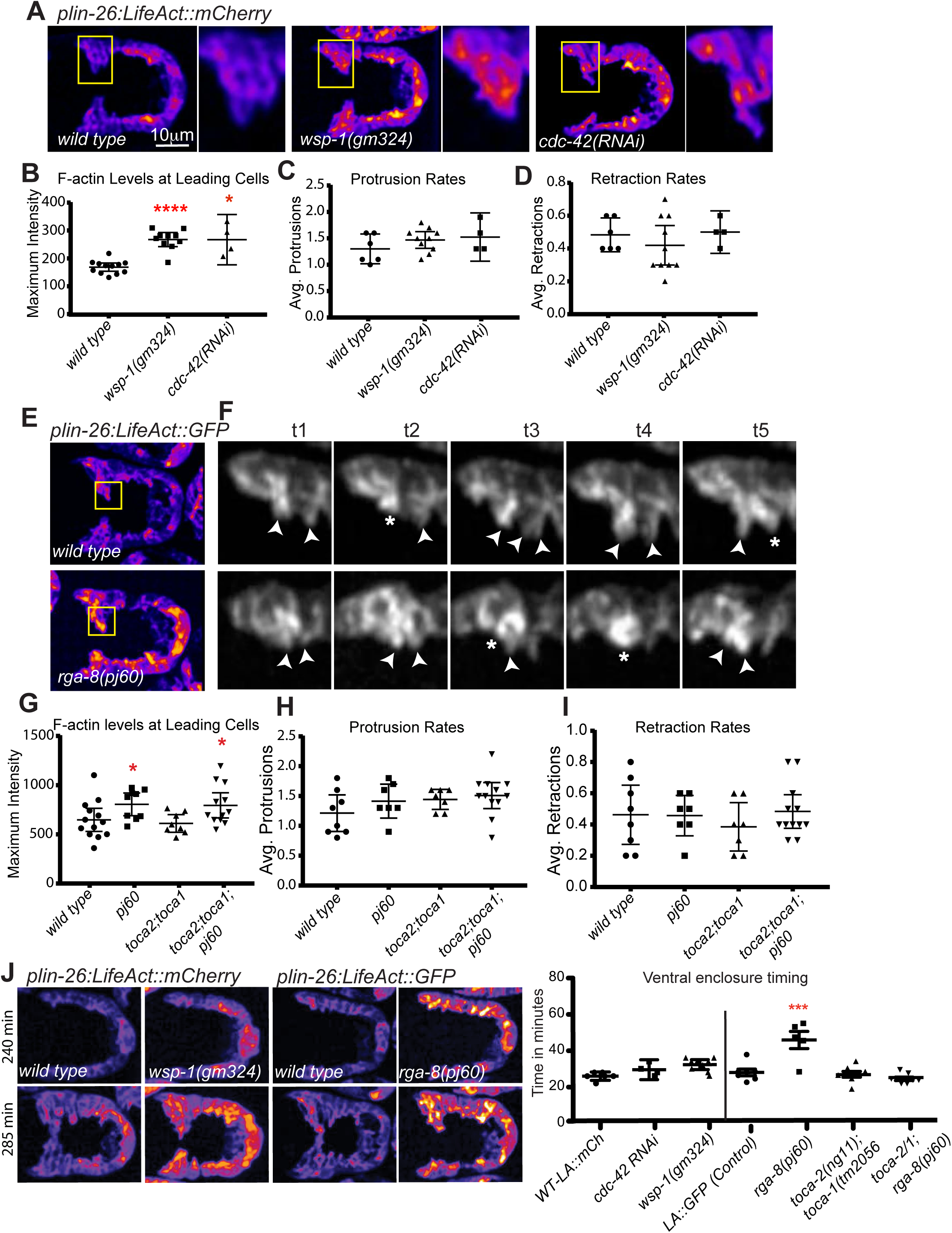
RGA-8 and WSP-1 alter levels of epidermal F-actin in migrating cells. (A & E) 270 minute embryos shown ventral up, surface focus, were used to compare levels of *plin-26::LifeAct::mCherry* (A) or *plin-26::LifeAct::gfp* (E) ([14]Gally et al., 2009, [36]Havrylenko et al., 2015) in migrating leading-edge epidermal cells during ventral enclosure. Both strains were used since some mutants mapped closely to the insertion sites of these integrated transgenes. Embryos were imaged 2 min intervals beginning at 240 min after first cleavage. Embryos were pseudocolored using the ‘Fire’ function in ImageJ, and intensity is shown from low (blue) to high (yellow). Yellow boxes enclose two leading cells, further amplified to the right. Levels (B,G): highest signal in the leading cells. Protrusions (C,H) and retractions (D,I) indicate average F-actin protrusions or retractions per time point in two leading cells during a 10 min period. F. In two Leading Cells, arrowheads mark protrusions and asterisks mark retractions. (J) These same movies were used to measure the timing of ventral enclosure, from the beginning of ventral-ward protrusions until the cells met at the midline. Quantitation of *plin-26::LifeAct::gfp* or *plin-26::LifeAct::mCherry* plotted for mean with 95% confidence interval. P Values ***= <0.001, ****= < 0.0001.

#### Protrusions dynamics are not significantly affected by the CDC-42 pathway

Ouellette and colleagues showed decreased dynamics in *wsp-1* mutants, measured as how much the leading-edge membrane is displaced in the leading cells ([21]Ouellette et al., 2016), although they did not examine F-actin levels. In contrast, we examined the dynamic formation of protrusion and retractions at the leading edge. We detected no significant change, though the number of protrusions was slightly increased in *wsp-(gm324)* and *rga-8(pj60)* mutants (Fig. 5C,D). Therefore, the elevated F-actin levels seem to not significantly perturb dynamics.

#### Speed of ventral enclosure is affected by the CDC-42 pathway

To test how changes in F-actin were affecting morphogenesis, we used the speed of migration as a phenotypic readout. To monitor timing for ventral enclosure using the epidermal F-actin movies, we measured from the time of the first protrusion to the first meeting at the ventral midline for the contralateral leading cells. *rga-8(pj60)* epidermal leading-edge cells migrate at slower rate (48min) compared to the wild type (24min), while *wsp-1(gm324)* and the *toca-2;toca-1* showed slight delays relative to controls that were not statistically significant.

### RGA-8 and CDC-42 pathway regulate NMY-2/myosin in epidermal cells

While CDC-42 has been connected to ventral enclosure, it is not clear which effectors of CDC-42 are responsible for this. CDC-42 regulates polarity (along with PAR-3/PAR-6/PKC) (Reviewed in [37]Goldstein & Macara 2007), cellular trafficking (through dynamin) (Reviewed in [38]Grant and Donaldson 2009), actin nucleation (through WSP-1 and formins) ([39]Lechler et al., 2001) and myosin contractility (through the myosin kinase MRCK-1) ([14]Gally et al., 2009). Since changes in the *cdc-42* pathway altered F-actin levels and slightly affected dynamics, we examined if RGA-8 is involved in NMY-2/myosin II regulation during ventral enclosure. First we crossed *mKate2::rga-8* with the *nmy-2::gfp* CRISPR allele ([40]Dickinson et al., 2013) and imaged the ventral epidermis during enclosure. There is some apparent colocalization along the ventral epidermis (Fig. 6A). Myosin is enriched as puncta at the front of migrating epidermal pocket cells that later converge at the ventral midline as the contralateral pocket cells meet [41,42] (green arrows in Fig. 6B). NMY-2/myosin II enrichment is highest in the ventral epidermis when the pocket cells meet, and then drops to basal levels as the embryo further elongates. To test for *cdc-42* pathway effects on this highly regulated myosin population, we measured levels of NMY-2/myosin II at first meeting of the contralateral pocket cells, as previously reported ([26]Wallace et al., 2018). *cdc-42* loss via RNAi caused variable changes in the level of NMY-2/myosin II in the migrating pocket epidermal cells, perhaps due to the complex effects of depleting *cdc-42*. In contrast, *wsp-1(gm324)* pocket cells consistently showed a 20% reduction compared to wild type. Similarly, the *rga-8(pj60)* putative null mutant caused a 22% reduction compared to wild type. Interestingly, removing both TOCA-1 and TOCA-2 did not significantly change NMY-2/myosin II enrichment, while a triple mutant that removes all three, *toca-2(ng11); toca-1(tm2016) rga-8(pj60)*, restored the level of NMY-2/myosin II back to wild type levels. A small deletion of *rga-8, ok3242*, predicted to truncate the C terminus, caused a 30% increase in myosin II/NMY-2, while the GAP point mutation, *rga-8*(*pj71)* caused a 15% drop, more similar to complete loss of *rga-8* (Fig. 6C). Measurements of myosin levels in the migrating leading cells showed similar changes, with approximately 20% drop in *rga-8(pj60)* (not shown). Thus, RGA-8 and CDC-42 pathway proteins regulate accumulation of myosin II/NMY-2 in enclosing epidermal cells (Fig. 6).

**Fig. 6.**
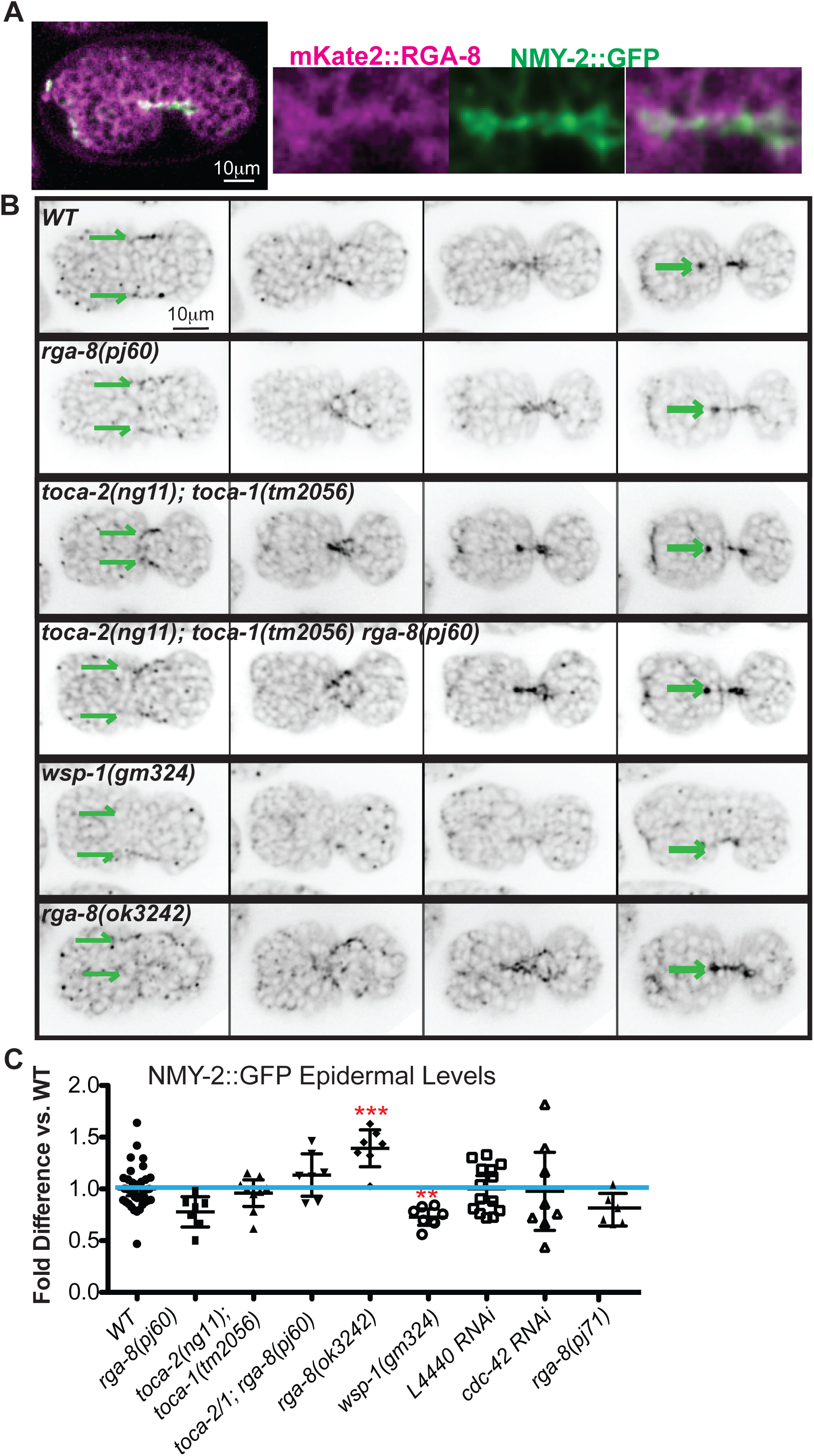
RGA-8 and WSP-1 regulate myosin in migrating epidermal cells. (A) 300 minute embryo expressing both NMY-2/myosin II CRISPR allele, *nmy-2::gfp* ([40]Dickinson et al., 2013), and *mKate2::rga-8(pj66)*, shown ventral up, and enlargement of the ventral midline during epidermal ventral enclosure. (B) Control and mutant strains are shown at 250, 280, 290 and 300 minutes, as the epidermal cells move toward and meet at the ventral midline. The signal from *plin-26::LifeAct::mCherry* was used to verify the cells shown are in the same plane as the epidermal cells. Parallel green arrows indicate the leading edge cells of the migrating epidermis. (C) *nmy-2::gfp* expression was measured when the contralateral pocket cells meet (approximately 300 min). Results are plotted as fold difference relative to controls. Plotted for mean with 95% confidence interval. Significance was calculated using 2 way ANOVA with Bonferroni’s post-test.

### RGA-8 and CDC-42 regulate the myosin light chain kinase, MRCK-1, in epithelia

To address how changes in *cdc-42* pathway genes may result in altered myosin levels, we focused on the myosin kinase MRCK-1. CDC-42 signaling directs MRCK-1 localization to activate myosin and cortical tension during *C. elegans* asymmetric cell division ([17]Kumfer et al., 2010), gastrulation ([43]Martson et al., 2016) and endocytic recycling ([44]Lant et al., 2015). We first tested if loss of *mrck-1* via RNAi affected *nmy-2::gfp* in the tissues studied here and detected a significant drop in *nmy-2::gfp* levels at the apical pharynx, and ventral epidermis (Fig. 7A). Next, we monitored MRCK-1 localization and levels using a CRISPR tagged strain, *mrck-1::gfp* ([43]Marston et al., 2016). The signal at the ventral epidermis also assembles in puncta, similar to *nmy-2::gfp*. Loss of *cdc-42* significantly reduced *mrck-1::gfp* enrichment at the apical pharynx during morphogenesis (Fig. 7B), and ventral epidermal signal was reduced by 15%, suggesting CDC-42 regulates MRCK-1 in these tissues. To test if RGA-8 regulates MRCK-1 in these tissues, we tested loss of *rga-8* via RNAi or the *pj71* GAP mutant and detected significantly increased *mrck-1::gfp* levels in both pharynx and ventral epidermis (Fig. 7B,C). We thus present evidence that a pathway involving the RhoGAP RGA-8 regulates activation of CDC-42, to regulate MRCK-1 that controls NMY-2 during epithelial morphogenesis.

**Fig. 7.**
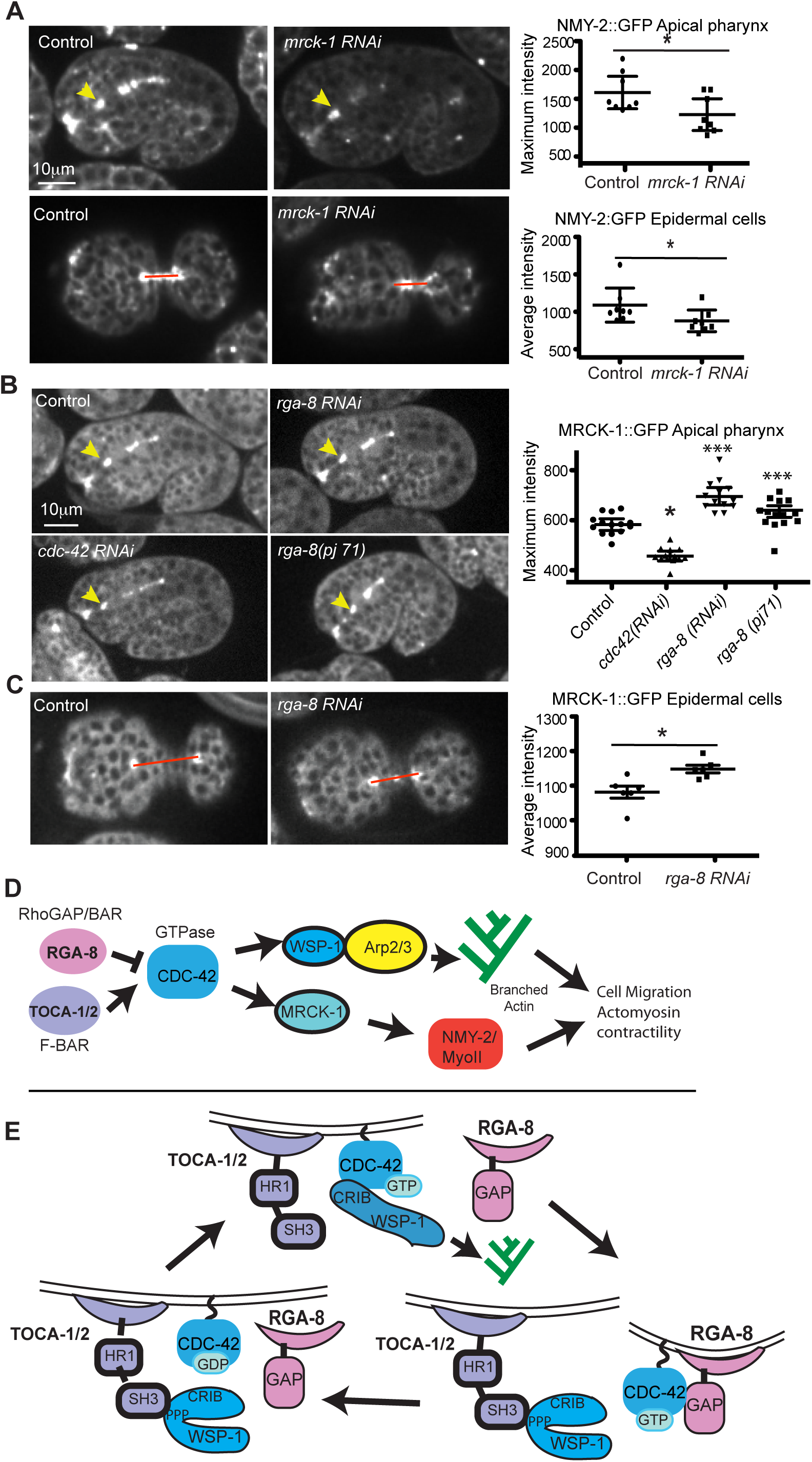
RGA-8 regulates the CDC-42 effector MRCK-1. (A) *mrck-1* RNAi effects on *nmy-2::gfp* in the apical pharynx, and ventral epidermis. *mrck-1::gfp* ([43]Marston et al., 2016) levels at the apical pharynx (B), and ventral epidermis during enclosure (C), in controls, *cdc-42 RNAi, rga-8 RNAi* and *rga-8(pj71)* embryos. n= at least 6 embryos for reach genotype. Significance was calculated using one-way ANOVA with Tukey post-test for B and by two-tailed student’s t test with Welch’s correction for A and C. P values, * = <0.05, *** = <0.001. (D) A genetic model for how the RhoGAP RGA-8 may work with TOCA-1 and TOCA-2 to regulate CDC-42 so that it can promote correct branched actin formation through WSP-1 and correct myosin activation through MRCK-1. This is a simplified model. For example, *mrck-1* is proposed to negatively regulate *mel-11*, which negatively regulates myosin activation ([14]Gally et al., 2009). (E) A molecular model, inspired by [50]Watson et al., 2016, to explain how a RhoGAP with BAR domain may work sequentially to TOCA-1/TOCA-2 F-BAR protein to support the cycle of CDC-42 activation that is needed during morphogenesis.

## Discussion

The GTPase activating protein (GAP) family consists of many members to regulate a smaller number of RHO-GTPases. In *C. elegans*, there are only 7 RHO-GTPases in comparison to 23 GAPs ([27]Neukomm et al., 2011). Numerous studies have contributed to our understanding of how different GAPs work on the RHO-family GTPases in *C. elegans* for early embryonic polarity establishment and cytokinesis ([17]Kumfer et al., 2010; [20]Beatty et al., 2013), clearance of corpses during cell death ([27]Neukomm et al., 2011), as well as epidermal morphogenesis ([21]Ouellette et al., 2016; [45]Zaidel-Bar et al., 2010). However, the function of the majority of the *C. elegans* GAP proteins is still unknown. This could be due to the mild phenotypes when individual GAPs are removed from the system. Some studies therefore use overexpression of these GAPs to investigate their function. Many *C. elegans* GAPs have no phenotypes on their own, so they are only examined in combination with loss of other proteins, for example, with hypomorphic alleles of the Cadherin component alpha catenin/*hmp-1* ([45]Zaidel-Bar et al., 2010). In disease studies, GAPs are often over-expressed, rather than missing. For example, high expression of homologs of RGA-8, including SH3BP1 (also named Nadrin and ARHGAP17) is associated with more invasive and chemo resistant cervical cancer ([46]Wang et al., 2018), and liver cancer ([47]Tao et al., 2016). These findings underscore the importance of careful study of the regulatory role of GAPs members, even if there is limited or mild phenotype in the null mutants.

In this study, we focused on the role of RhoGAP RGA-8, a proposed new regulator of CDC-42, during epidermal morphogenesis, and addressed two key questions – (1) how does it affect epidermal morphogenesis; (2) which CDC-42-dependent events does this proposed GAP protein regulate? We found evidence that RGA-8 regulates active CDC-42, and that it functions together with CDC-42 to regulate ventral enclosure by regulating the level of actin and non-muscle myosin (NMY-2) in the migrating embryonic epidermal cells. These changes correlate with effects on the speed of morphogenetic events. We further showed that RGA-8 alters apical/basal distribution of active CDC-42 in two other epithelia, the pharynx and intestine. We propose that the RhoGAP RGA-8 regulates morphogenesis through effects on CDC-42, based on genetic interactions, expression patterns, and phenotypes.

### Genetic interactions – embryonic lethality

*cdc-42 RNAi* led to embryonic lethality due to failures in morphogenesis and this was significantly reduced by the *rga-8* deletion allele, *pj60. rga-8(pj60)* resulted in additive effects with the *toca-2(ng11);toca-1(tm2056)* double mutant. TOCAs are known interactors of CDC-42, that regulate branched actin by activating WSP-1/WASP. Interestingly, *wsp-1(gm324); rga-8(pj60)* double mutants resulted in intermediate embryonic lethality closer to that of *wsp-1(gm324)* mutants, which suggested *wsp-1* is epistatic to *rga-8*. These genetic interactions suggested a pathway where TOCA-1 /TOCA-2 and RGA-8 mutually regulate CDC-42, to ensure appropriate activation of WSP-1 (Table 1, 2).

### Expression patterns

A CRISPR tagged mKate2::RGA-8 strain showed that while RGA-8 is broadly and diffusely expressed, the highest expression was in regions of epithelia where GFP::CDC-42 is also enriched, including apical pharynx and intestine. This enrichment begins in embryos as these tissues polarize and mature, and continues into adulthood (Fig. 2). By comparison, active CDC-42, as shown by the *Pcdc-42::gbd-wsp-1::gfp* CDC-42 biosensor, has a distinct pattern in the developing intestine (Fig. 3). Epidermal, pharyngeal and intestinal distribution of *Pcdc-42::gbd-wsp-1::gfp* depend on RGA-8 (Fig. 4). Despite showing changes in apical/basal membrane distribution and levels of *gbd-wsp-1::gfp*, most null *rga-8* embryos are viable, suggesting the effects of RGA-8 on CDC-42 are redundant with other proteins, perhaps other GAPs. Since RGA-8 has a BAR domain, we investigated if it localized to specific subcellular membranes. We found it has diffused enrichment around apical epithelia, and may only be transiently enriched at epidermal membranes (Fig. 3).

### Shared phenotypes with CDC-42 pathway

*T*he phenotypes of three *rga-8* alleles and of *cdc-42* pathway components suggested a common function during epithelial morphogenesis. In particular, we found that deletion alleles of the *toca-2; toca-1* double, or *rga-8*, or *wsp-1*, or *cdc-42* RNAi, led to similar embryonic lethality with arrests during both ventral enclosure and elongation, and expanded intestinal apical lumen (Fig. 1B,C). Mutations in RGA-8 led to highly penetrant changes in epidermal cell migration timing, in levels of epidermal F-actin and in levels of epidermal myosin (Fig. 5, 6). Similar changes were also seen in mutations of *toca-2;toca-1, wsp-1*, and *cdc-42 RNAi*. Embryonic lethality was not fully penetrant for any of the RGA-8 alleles. This is also true for complete loss of *toca-2; toca-1* (13%) or *wsp-1* (35%). One explanation for this ability of embryos to sometimes survive without the contribution of the TOCAs/RGA-8/CDC-42/WSP-1 pathway is that this pathway is responsible for only part of the events of epidermal morphogenesis. In contrast, the CED-10/WAVE pathway leads to 100% embryonic lethality, suggesting that Rac/WAVE regulation of Arp2/3 cannot be compensated for by other pathways. CDC-42 has additional proposed roles besides regulating branched actin through WSP-1, including regulation of formins, and thus linear actin, and regulation of Myosin through two kinases, PAK-1 and MRCK-1 (Gally et al., 2009). Thus, CDC-42’s full contribution to epidermal morphogenesis will require analysis of additional regulatory pathways.

### Actin phenotypes

A well-studied function of CDC-42 is to regulate branched actin dynamics via the WASP/WSP-1 protein. Reduction of *cdc-42* by RNAi under the conditions used here caused some embryos to fail elongation, and some embryos to fail ventral enclosure. Similar defects were seen for the *wsp-1* deletion, *gm324*. If we consider that *wsp-1(gm324)* is a read out for defective CDC-42-dependent branched actin, then its phenotypes of elevated levels of F-actin and decreased myosin during ventral enclosure suggest CDC-42/WSP-1 dependent branched actin normally interacts with and promotes myosin to support the migrations. The elevated F-actin seen could be default linear F-actin, since studies in yeast suggest that in the absence of branched actin nucleators, formins take over ([48]Suarez and Kovar, 2016).

The *toca-2; toca-1* double mutant did not significantly alter F-actin or myosin levels. However, *toca-2; toca-1* mutants are defective in ventral enclosure and elongation ([23]Giuliani et al., 2009). This may indicate TOCAs play a distinct role regulating CDC-42 compared to RGA-8. For example, TOCA-1/-2 and CDC-42 are proposed to promote trafficking, and the morphogenetic effects could be linked to their trafficking roles ([49]Fricke et al., 2009). TOCA-2/TOCA-1 and RGA-8 may also control different populations of F-actin. While TOCA-2/TOCA-1 is thought to regulate branched actin formation through WASP/WSP-1, TOCA-2/TOCA-1 can also bind two components of the WAVE/SCAR complex, WVE-1 and ABI-1 ([23]Giuliani et al., 2009), suggesting potential crosstalk between the WAVE/SCAR pathway and TOCAs to support epidermal migrations.

### Myosin phenotypes

Our analysis of myosin behavior in epidermal cells suggested that myosin plays a role during epidermal ventral enclosure (this study and [26]Wallace et al., 2018). Interestingly, Oulette and colleagues reported that overexpression of CDC-42 caused a decreased protrusion rate of leading-edge cell migration, resulting in delayed ventral enclosure (Ouellette et al., 2016). Similarly, we found that loss of *rga-8*, which as a GAP might result in excess active CDC-42, resulted in delayed leading-edge cell migration during ventral enclosure. This suggests that myosin activity during epidermal ventral enclosure may be under the control of CDC-42, and RGA-8. In support of this, *rga-8* loss via RNAi led to increased *mrck-1::gfp* (Fig. 7), and decreased *nmy-2::gfp* (Fig. 6), suggesting misregulated CDC-42 results in misregulated MRCK-1 that alters levels of myosin, and also, perhaps its localization. Higher resolution live imaging of myosin during these dynamics events can address this model.

### Conclusion

Analysis of some of the 23 *C. elegans* Rho GAPs has broadened our view of how the three main Rho GTPases, Rac, Rho and CDC-42, contribute to morphogenesis. Our analysis of Rho GAP HUM-7/Myo9 showed it affects Rho signaling, and is needed to attenuate Rho in the migrating epidermal cells, to prevent excessive myosin and protrusions ([26]Wallace et al, 2018). The Nance lab showed that RhoGAP PAC-1 supports elongation by regulating CDC-42 ([24]Zilberman et al., 2007), while the Jenna lab showed that RhoGAP RGA-7 plays a role possibly through regulation of CDC-42 ([21]Ouellette et al., 2016). Here we add the analysis of an intriguing RhoGAP, RGA-8, with both RhoGAP and BAR domains. By extension of the proposed role for TOCAs in bringing together WSP-1 and CDC-42 to promote active CDC-42-GTP ([50]Watson et al., 2016), we propose RGA-8 may use its GAP domain to bind to active CDC-42-GTP, and generate CDC-42-GDP and renewal of the CDC-42 cycle (Fig. 7E). It is thought that GAP hydrolysis of GTPases is the essential regulatory step for GTPase function. For CDC-42, studies in budding and fission yeast have shown that localized activation by GEFs is not enough to generate locally restricted CDC-42 activity, and GAPs are needed to prevent spread of the active form (Reviewed in [51]Chiou et al., 2017). In fission yeast, the GTPase Ras1 requires inactivation by its GAP, or it suffers loss of spatial information ([52]Merlini et al., 2018). Models have been proposed to explain how GAP activity can promote GTPase activity, even when the GAP and GEF for a GTPase colocalize. For example, in the one-cell embryo of *C. elegans* the levels of the GTPase RHO-1/RhoA, are modulated by its GEF ECT-2 and its GAPs RGA-3 and RGA-4, which colocalize at the anterior domain ([19]Schonegg et al., 2007). RGA-8 has a BAR domain, which suggests it can tubulate membranes (Reviewed in [53]Carman and Rodriguez, 2018). Our results show enrichment of the RhoGAP RGA-8 at apical regions, and RGA-8 regulation of a CDC-42 biosensor with distinct apical/basal distribution in epithelia, supporting that this GAP plays an important role fine-tuning the localization of active CDC-42 as organs and tissues polarize. Further experiments will need to investigate which of the many cellular processes regulated by CDC-42 are promoted by the RhoGAP-BAR protein RGA-8.

## MATERIALS AND METHODS

### Strains

All strains used in this study are listed in Table 3. Strains were either received from the CGC (Caenorhabditis Genetics Center, USA), the NBRP (National Bio-Resource Project, Japan), or individual labs listed below, or were generated for this study. All strains were grown at 23°C unless otherwise stated.

**Table 3.** Strains used in this study –.

| Strain Name | Genotype | Source |
| --- | --- | --- |
| N2 | *Wild type* | CGC |
| NG0324 | *wsp-1(gm324)* | CGC via [11] |
| MT5013 | *ced-10(n1993)* | CGC |
|  | *toca-2(ng11); toca-1(tm2056)* | [23]Giuliani et al., 2009 |
| OX680 | *rga-8(pj60)* | This study |
| OX808 | *rga-8(pj71)* | This study |
| OX768 | *wsp-1(gm324); rga-8(pj60)* | This study |
| OX711 | *ced-10(n1993); rga-8(pj60)* | This study |
| OX794 | *let-502(sb118ts); rga-8(pj60)* | This study |
| OX748 | *toca-2(ng11); toca-1(tm2056) rga-8(pj60)* | This study |
| RB2384 | *rga-8(ok3242)* | CGC (& outcrossed) |
| OR113 | *dlg-1::gfp; rol-6(su1006)* | Chris Rongo |
| OX725 | *dlg-1::gfp; rol-6(su1006); rga-8(pj60)* | This study |
| OX785 | *dlg-1::gfp; rol-6(su1006); rga-8(ok3242)* | This study |
| OX749 | *dlg-1::gfp; rol-6(su1006); toca-2(ng11); toca-1(tm2056) rga-8(pj60)* | This study |
| OX385 | *dlg-1::gfp; rol-6(su1006); toca-2(ng11); toca-1(tm2056)* | Soto Lab; [23]Giuliani et al., 2009 |
| OX889 | *dlg-1::gfp; rol-6(su1006); rga-8(pj71)* | This study |
| OX684 | *rga-8(pj66[mKate2::FLAG:: rga-8 + LoxP])* | This study |
| VJ402 | *fgEx11 (erm-1::gfp, rol-6)* | Verena Gobel, [33]Gobel et al., 2004 |
| LP169 | *hmp-1(cp20[hmp-1::gfp + LoxP unc-119(+)] LoxP))* | [43]Marston et al., 2016 |
| OX724 | *rga-8(pj66[mKate2::FLAG:: rga-8 + LoxP]); dlg-1::gfp; rol-6(su1006)* | This study |
| OX734 | *rga-8(pj66[mKate2::FLAG:: rga-8 + LoxP]); erm-1::gfp* | This study |
| OX726 | *rga-8(pj66[mKate2::FLAG:: rga-8 + LoxP]); hmp-1::gfp* | This study |
| OX721 | *rga-8(pj66[mKate2::FLAG:: rga-8 + LoxP]); arp-2::gfp* | This study |
| OX747 | *rga-8(pj66[mKate2::FLAG:: rga-8 + LoxP]); nmy-2::gfp* | This study |
| WS4700 | *opls295 [cdc-42p::GFP::cdc-42(genomic)::cdc-42 3'UTR + unc-119(+)]* | CGC, [32]Neukomm et al., 2014 |
| OX880 | *opls295 [gfp::cdc-42]; rga-8(pj60)* | This study |
| FT1459 | *unc-119(ed3); xnls506 [cdc42P::gst-gfp-wsp-1gbd, unc-119(+)]* | [24]Zilberman et al., 2017 |
| OX762 | *rga-8(pj66[mKate2::FLAG:: rga-8 + LoxP]); opls295 [cdc-42p::GFP::cdc-42(genomic)::cdc-42 3'UTR + unc-119(+)]* | This study |
| OX832 | *rga-8(pj66[mKate2::FLAG:: rga-8 + LoxP]); unc-119(ed3); xnls506 [cdc42P::gst-gfp-wsp-1gbd, unc-119(+)]* | This study |
| JUP30 | *unc119(ed3); ls[Plin-26::Lifeact::GFP::unc54 3'UTR; Cb-unc119]* | [36]Havrylenko et al., 2015 |
| LP162 | *nmy-2(cp13[nmy-2::GFP + LoxP]) I.* | [40]Dickinson et al., 2013 |
| OX732 | *nmy-2::gfp; rga-8(pj60)* | This study |
| OX846 | *nmy-2::gfp; toca-2(ng11); toca-1(tm2056)* | This study |
| OX855 | *nmy-2::gfp; toca-2(ng11); toca-1(tm2056) rga-8(pj60)* | This study |
| OX747 | *nmy-2::gfp; rga-8(pj66 [mKate2::rga-8])* | This study |
| OX756 | *nmy-2::gfp; Plin-26::LifeAct::mCherry* | This study |
| OX833 | *nmy-2::gfp; wsp-1(gm324); plin-Lifeact-mCherry* | This study |

### Strain generation using CRISPR

CRISPR strains were made using the SEC strategy ([31]Dickinson et al., 2015). PAM sites were identified with the help of http://crispr.mit.edu/ website from MIT as a guide. Generation of guide RNA (gRNA) plasmids was made using either sewing PCR method with the primers MSo1204-1205, MSo1304-1305 and MSo1367-1340 ([54]Kim et al., 2014), or digestion and ligation using the primers MSo1585-1586 ([55]Arribere et al., 2014). To test Cas9 cutting and the gRNAs we used co-CRISPR ([54]Kim et al, 2014). *To generate tagged genes*, we used the plasmid rescue method from the Goldstein lab (Dickinson et al., 2015). Briefly, the 5’ flanking arms were generated using primers MSo1402-1403 and the 3’ flanking arms were generated using primers MSo1404-1405. Both arms were cloned into the pDD285 vector using Gibson assembly kit from NEB Cat# E2611. *To generate deletion mutants*, we used co-CRISPR method by the Fire lab to select for worms that have a successful Cas9 cutting event ([55]Arribere et al., 2014). *To generate specific point mutants*, verified sgRNA and rescue oligos containing the desired point mutation as well as mutated PAM site were co-injected with *dpy-10* sgRNA. Silent mutation that inserted a new AvaI restriction enzyme site was engineered on the rescue oligos (MSo1602) to ease the screening process of identifying the mutants. DNA sequencing was done to verify the mutations and inserts for all strains. All strains were back crossed to wild type at least three times to minimize possible off-target effects by the gRNAs and Cas9. Primers used for strain generation are listed in Table 4.

**Table 4.**
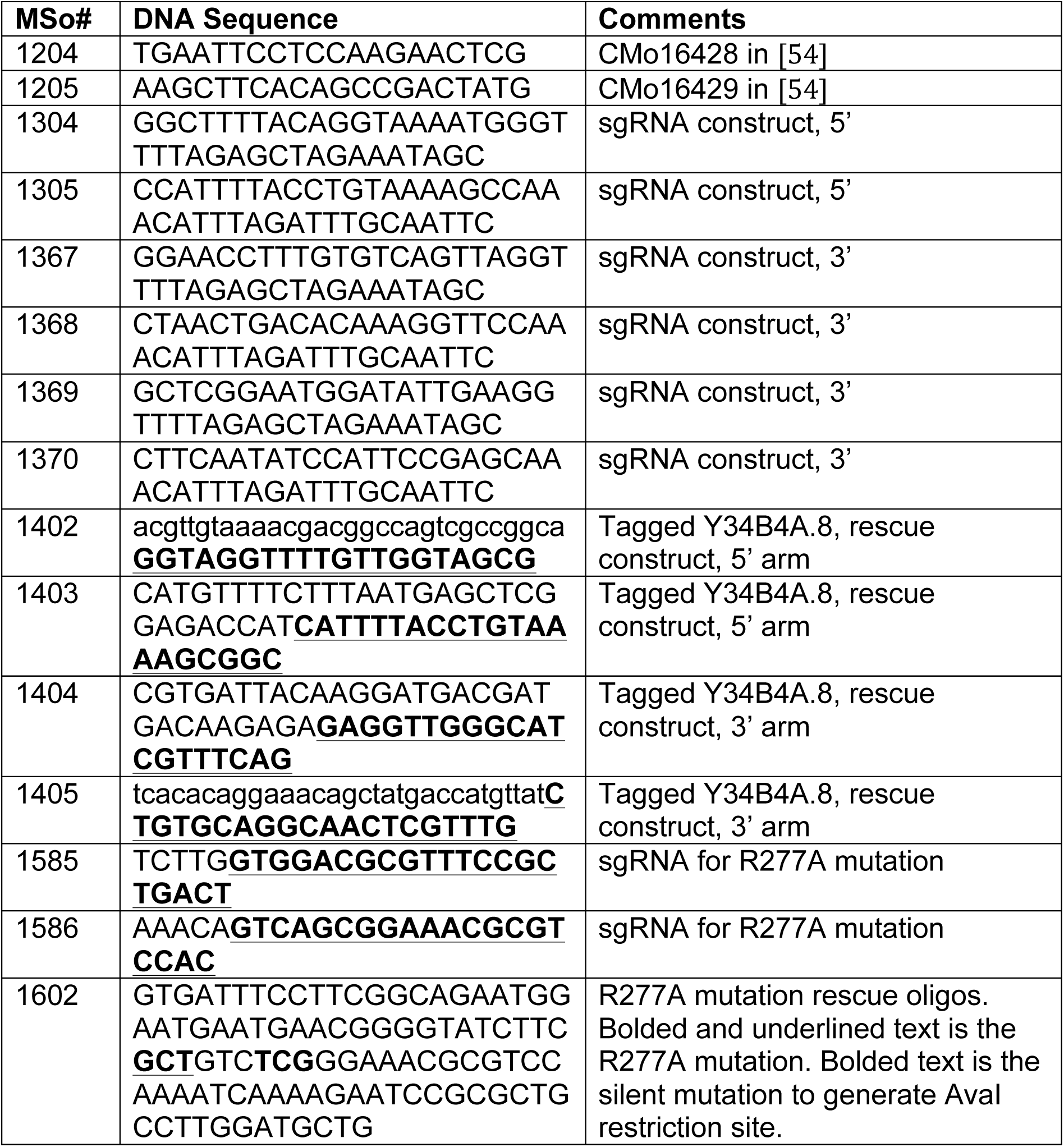
DNA primers used to generate RGA-8 CRISPR alleles.

### RNAi experiments

All RNAi bacterial strains used in this study were administered by the feeding protocol as in Bernadskaya et al. ([35]2012). Most RNAi clones were obtained from the Driscoll lab library at Rutgers University or generated in the lab. RNAi feeding experiments were done at 23°C unless otherwise mentioned. Worms were synchronized and transferred onto seeded plate containing RNAi-expressing bacteria. Embryos were scored for lethality at either 48 hours (for *mel-11* and *cdc-42* RNAi) or 72 hours (for all other genes).

### Live Imaging

For all live imaging shown, embryos at the 2 to 4 cell stage were dissected from adult hermaphrodites and mounted onto 3% agarose pads, covered with #1.5 cover slip, and sealed with Vaseline. Embryos were then incubated at 23°C for 240 minutes. Imaging was done in a temperature-controlled room set to 23°C on a Laser Spinning Disk Confocal Microscope with a Yokogawa scan head, on a Zeiss AxioImager Z1m Microscope using the Plan-Apo 63X/1.4NA or Plan-Apo 40X/1.3NA oil lenses. Images were captured on a Photometrics Evolve 512 EMCCD Camera using MetaMorph software, and analyzed using ImageJ. Some images (Fig. 1) were done on a Zeiss Axioskop 2 microscope, using Plan-Apo 40X/1.3NA oil lens, captured with a Roper camera. Controls and mutants were imaged within 3 days of each other with the same imaging conditions. All measurements were performed on raw data. For fluorescent measurements, background intensity was subtracted by using a box or line of the same size and measuring average intensity in the same focal plane, near the embryo. Actin Intensity measurements, protrusion and retraction analysis as previously done ([26]Wallace et al., 2018) on embryos imaged at 2-minute intervals for at least 120 minutes beginning at 240 min after 2 to 4 cell stage.

### Myosin measurements

To compare myosin levels as pocket cells meet, a rectangular box enclosing the pocket cells as they first touch (approximately 320min) was drawn (yellow box in Fig. 6C), Time points shown and measured are those when the epidermal cells first touch based on epidermal F-actin signal. To measure myosin puncta on the same focal plane as the epidermis, we co-localized the highest Plin-26::LIFEACT::mCherry intensity with NMY-2::GFP and recorded the highest of three measurements per embryo (Fig. 6B). Maximum intensity values were recorded after subtracting the average background fluorescence. The graph in Fig. 6B records the relative level of NMY-2::GFP, after normalization of either WT, or WT on control RNAi, to 1.

### Statistical Analysis

For grouped data statistical significance was established by performing a two-way Analysis of Variance (ANOVA) followed by the Bonferroni multiple comparison post-test. For ungrouped data an unpaired t-test, the unequal variance (Welch) t test, was used. Error bars show 95% confidence intervals. Asterisks (*) denote p values *= p<.05, ** = p<0.001, *** = p<0.0001, ***=p<0.00001. All statistical analysis was performed using GraphPad Prism.

## Acknowledgements

We thank the NCRR-funded *Caenorhabditis* Genetics Center (CGC), funded by NIH Office of Research Infrastructure Programs (P40 OD010440), the National Bio-Resource Project, MEXT, Japan, Paul Mains, Verena Gobel and Bob Goldstein for strains, and the Goldstein and Mello labs for advice on CRISPR. We thank Soto lab members for comments on this manuscript. We thank the National Institutes of Health Shared Instrumentation Program (S10OD010572) for the Spinning Disk used in this research.

## Funding

This work was supported by the National Institutes of Health [R01GM081670 to M.C.S. and an Institutional Research and Academic Career Development Award (K12GM093854) to A.W.]; the RWJMS Graduate Office [6 month thesis completion grant to H.R.]; and NJCRR, the New Jersey Commission for Cancer Research [DFHS18PPC044 to S.S.].

